# The olfactory co-receptor IR8a governs larval-frass mediated competition avoidance in a hawkmoth

**DOI:** 10.1101/725820

**Authors:** Jin Zhang, Sonja Bisch-Knaden, Richard A. Fandino, Shuwei Yan, George F. Obiero, Ewald Grosse-Wilde, Bill S. Hansson, Markus Knaden

## Abstract

Finding a suitable oviposition site is a challenging task for a gravid female moth. At the same time, it is of paramount importance considering the limited capability of most caterpillars to relocate to alternative host plants. The hawkmoth, *Manduca sexta* (Sphingidae), oviposits on solanaceous plants. Larvae hatching on a plant that is already attacked by conspecific caterpillars can face food competition, as well as an increased exposure to predators and induced plant defenses. Here, we show that frass from conspecific caterpillars is sufficient to deter a female *M. sexta* from ovipositing on a plant and that this deterrence is based on the frass-emitted carboxylic acids 3-methylpentanoic acid and hexanoic acid. Using a combination of genome editing (CRISPR/Cas9), electrophysiological recordings, calcium imaging and behavioral analyses we demonstrate that the ionotropic co-receptor IR8a is essential for acid-mediated frass avoidance in ovipositing hawkmoths.

## Introduction

For insects, finding appropriate sites for oviposition is a challenging task and the decision of a gravid female will have clear consequences for the fitness of its progeny. Due to fragility and limited mobility, the offspring faces many threats: limited food availability, intra- and interspecific competition, predation and attack by parasitoids. Therefore, gravid females must carefully examine the environment prior to selecting the oviposition site. For this, they utilize visual^1, 2^, gustatory^3, 4^, mechanosensory^5^, as well as olfactory^6, 7^ cues. Among these modalities, olfaction plays a pivotal role in an insect’s life, as it provides information not only about oviposition sites but also about other biologically relevant resources such as food and mating partners^8^.

Insects rely on a sophisticated olfactory system to detect volatile chemicals in the environment. Several protein families are involved, with odorant receptors (ORs) and ionotropic receptors (IRs), two types of ligand-gated ion channels, being the key detecting elements^9-11^. On the surface of the antenna, the main olfactory organ, numerous hair-like structures (sensilla) contain olfactory sensory neurons (OSNs), which represent the basic units of sensory perception. Sensilla involved in olfaction occur in three morphological types: basiconic, trichoid, and coeloconic. In the vinegar fly, *Drosophila melanogaster*^9, 12^, as well as in other investigated insect species^13-15^, ORs are expressed in the dendritic membrane of OSNs housed in basiconic and trichoid sensilla, whereas IRs are expressed by OSNs housed in coeloconic sensilla. ORs are extremely divergent and different insect species express from ten *OR* genes in head lice^16^ to more than 300 in ants^17^. The OR type expressed in an OSN dictates the odorant specificity of the neuron^18^. ORs are co-expressed together with the conserved odorant receptor-co-receptor (Orco), which is essential for dendritic localization of ORs and OR-dependent odorant detection^10, 19^. IRs usually are less divergent^20^ and at least two IR co-receptors, IR8a and IR25a, form ligand-gated ion channels with other odorant-tuned IRs^9, 21^. The different receptor types, however, do not only differ regarding their local expression but also in their response profiles. While most ORs are broadly tuned to alcohols, aldehydes, aromatics, esters, or terpenes^18^, IRs primarily respond to a restricted subset of odors including mainly acids and amines^22^. At least in *Drosophila* and *Aedes aegypti*, IR8a is required for acid detection^23, 24^. IR25a, on the other hand, seems to be co-expressed with IRs responding to amines^25^ and is also involved in the detection of temperature^26^, humidity^27^ and salt^28^.

The tobacco hawkmoth *Manduca sexta* (Lepidoptera: Sphingidae) is an established model for insect olfaction^13^ and odor-guided behavior^29^. The recent identification of 73 *OR* genes and 21 olfactory *IR* genes and their expression patterns in male and female moths^13^ and the establishment of the Crispr/Cas 9 technique in *M. sexta*^30^ has made the species to an even more powerful model for olfactory neuroethology. The larvae of these moths feed on various plants of the family Solanaceae, including coyote tobacco (*Nicotiana attenuata*) and jimson weed (*Datura wrightii*). It was reported that a single *M. sexta* caterpillar consumes 1-10 tobacco plants until pupation^31^, resulting in complete defoliation of the plants and accumulation of frass under the plant (Fig. 1 a). Therefore, it is crucial for female *M. sexta* to find a suitable host plant that is not already occupied by a conspecific larva.

**Figure 1.**
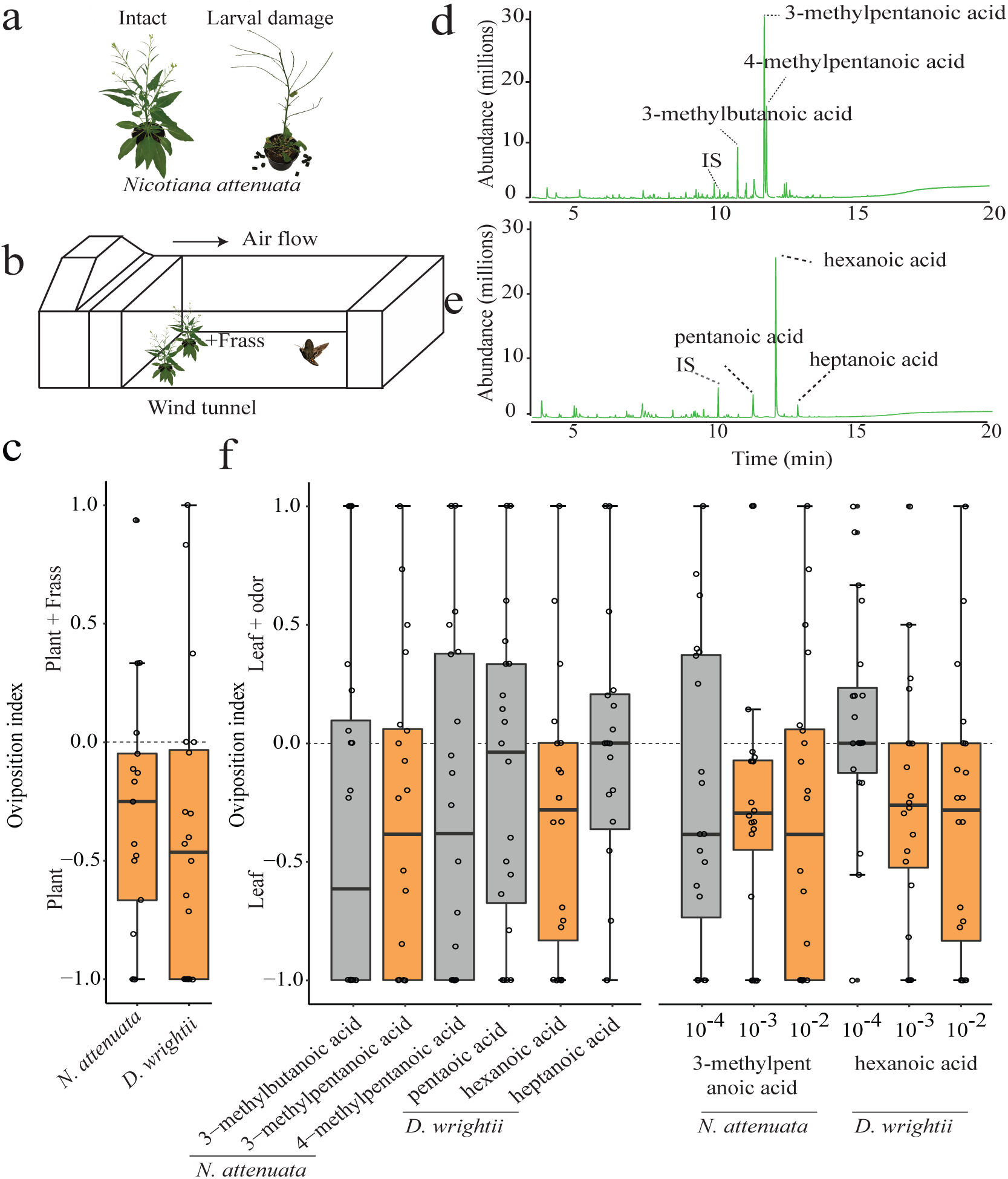
*M. sexta* oviposition on *N. attenuata* and *D. wrightii* are affected by larval frass. (**a**) *N. attenuata* plant with and without larval damage. (**b**) Schematic drawing of wind tunnel assay. (**c**) Oviposition index of mated females toward frass of *M. sexta* caterpillars reared on *N. attenuata* and *D. wrightii*. Oviposition index = (number of eggs on plant with frass - number of eggs on plant without frass) / total egg number. (**d**) GC-MS profile of headspace of frass from *M. sexta* caterpillar reared on *N. attenuata*. IS, internal standard. (**e**) GC-MS profile of headspace of frass from *M. sexta* caterpillar reared on *D. wrightii*. IS, internal standard. (f) Oviposition index of gravid females to carboxylic acids emitted from *M. sexta* caterpillar frass (left panel). (**f**) Oviposition index of gravid females to various doses of 3-methylpentanoic acid and hexanoic acid (right panel). Deviation of the index against zero was tested with Wilcoxon signed-rank test (n=17-20). * p<0.05. Boxplots depict median and upper and lower quartile; whiskers depict quartiles +/− 1.5× the interquartile range (IQR). Any data points above the superior or below the inferior whisker values are considered as outliers. All data were included in the statistical analysis. Orange boxes depict significant repulsion of frass and/or individual compounds.

Volatiles emitted from larval frass have been shown to act as kairomones and attract parasitoids and predators^32-35^. The smell of larval frass, therefore, not only indicates the occupancy of the host plant, and the resulting potential for intra-specific competition, but also an increased susceptibility to parasitization and predation. Hence, female moths should avoid sites that are already occupied by conspecific larvae and could do so by e.g. detecting chemical cues emanating from larval frass. In several insect species, female oviposition has been found to be deterred by conspecific larval frass^36^. Thus, larval frass alone is sufficient to signal potential competition to the female. However, the molecular and cellular mechanisms by which female insects avoid frass remain unknown.

Here we investigate whether the oviposition of *M. sexta* is deterred by frass from its larvae. We first show that *M. sexta* females, like other insects, display oviposition aversion toward conspecific caterpillar frass stemming from different host plants. Next, we identify specific carboxylic acids emitted from the frass as key compounds that confer oviposition aversion. By performing electrophysiological recordings, calcium imaging and behavioral analyses with mutant moths that either lack *Orco*, or one of the IR co-receptors, *Ir8a* or *Ir25a*, we demonstrate that IR8a is essential for acid-mediated frass avoidance during oviposition.

## Results and discussion

### Frass of caterpillars fed on *N. attenuata* repels oviposition

To test whether gravid females of *M. sexta* avoid ovipositing in the presence of frass, we tested their behavior in a two-choice assay in a wind tunnel. The moths were allowed to oviposit for 3 min either on an undamaged *N. attenuata* plant that was equipped with 10 g of larval frass (from caterpillars which had fed on other *N. attenuata* plants) or on an undamaged control plant (Fig. 1 b). In these experiments, the moths laid on average 14.3 ± 1.7 eggs (mean ± SEM) during the 3 min test on both plant. The allocation of eggs depended on the presence of frass with the moths laying significantly less eggs on the plant with caterpillar frass as compared to the control plant (Fig. 1 c). A similar preference was observed when moths were given a choice between a plant with frass and a control plant without frass in a steady-air tent (Supplementary Fig. 1), confirming that frass avoidance is consistent in different behavioral paradigms. We conclude that even in the absence of plant damage caterpillar frass induces oviposition avoidance in *M. sexta*. Former studies suggested that ovipositing *M. sexta* females mainly use plant- and larva-derived odors to avoid competition^37, 38^. In our study, where the amount of frass was higher (but still ecologically reasonable, as we used frass that was produced by a single larvae during one night), frass alone was sufficient to induce oviposition avoidance. Females tested in our experiments were raised on artificial diet and had no prior experience with the plants or the frass. We therefore conclude that the frass-induced oviposition avoidance is innate.

### 3-methylpentanoic acid and hexanoic acid govern oviposition avoidance to larval frass

To identify the active compound responsible for frass avoidance, we raised *M. sexta* caterpillars on *N. attenuata* plants and afterwards collected the headspace of the resulting frass using a solid phase micro extraction (SPME) fiber. Gas chromatography–mass spectrometry (GC-MS) analysis revealed similar results as in a previous study^33^, with 3-methylpentanoic acid being the most abundant compound followed by two other branched aliphatic acids, 3-methylbutanoic acid and 4-methylpentanoic acid (Fig. 1 d).

To investigate the impact of these compounds on *M. sexta* oviposition, we pipetted 10 µl of one of the compounds (diluted 10^−2^ in mineral oil) on a filter paper and attached this filter paper 2 cm upwind of a detached *N. attenuata* leaf before presenting this leaf to a mated female in the wind tunnel. When compared to a control leaf, where the attached filter paper just contained the solvent, only 3-methylpentanoic acid elicited significant avoidance (Fig. 1 f, left panel). To address the behavioral sensitivity of *M. sexta* toward 3-methylpentanoic acid, we further performed wind tunnel test with lower amounts of the compound and identified the behavioral threshold to be between 10^−4^ and 10^−3^ dilutions (i.e. between 9.3 µg and 93 µg) (Fig. 1 f, right panel).

Having shown that frass from caterpillars reduces the attraction of *N. attenuata* plants to ovipositing females, we asked whether this holds true also for the relationship of *M. sexta* and its other main host plant *D. wrightii.* We now let female moths choose to oviposit either on a *D. wrightii* leaf that was equipped with frass (from caterpillars raised on *D. wrightii*) or on a control leaf without frass. Again, females preferred to oviposit on the control leaf (Fig. 1 c), suggesting that also at the host plant *D. wrightii*, frass induces oviposition avoidance in *M. sexta* females.

To identify the active compounds responsible for frass avoidance in *D. wrightii*, we raised *M. sexta* caterpillars on *D. wrightii* plants and then collected and analyzed the volatiles as before. This time the chemical profile of the frass was dominated by hexanoic acid, and accompanied by two other minor compounds, heptanoic acid and pentanoic acid (Fig. 1 e). When again an ovipositing female had to choose between a *D. wrightii* leaf that was equipped with one of the three acids and a control leaf, only leaves with hexanoic acid (10 µl at 10^−2^ dilution) were avoided (Fig. 1 f, left panel). This avoidance could still be observed even when we reduced the amount of hexanoic acid tenfold (Fig. 1 f, right panel). We conclude that hexanoic acid is the major compound governing frass avoidance of ovipositing females in the context of *D. wrightii.* When performing choice experiments in the wind tunnel with additional aliphatic acids and the two host plants, we found that only six-carbon aliphatic acids elicited avoidance (Supplementary Fig. 2).

### Both the odorant co-receptor Orco and the ionotropic co-receptor 8a participate in acid sensing

To determine which olfactory pathway is governing the detection of 3-methylpentanoic acid and hexanoic acid, we performed electroantennography (EAG) measurements on wild type (WT) moths, and on odorant co-receptor heterozygous (*Orco*+/−) and homozygous (*Orco*−/−) moths that were recently generated in our lab using CRISPR/Cas9 genome editing^30^. While WT moths and *Orco*+/− moths exhibited robust EAG responses to the acids, *Orco*−/− moths showed reduced responses (Supplementary Fig. 3). However, clear EAG responses to the acids remained, indicating that the IR pathway is also involved in acid detection.

**Figure 2.**
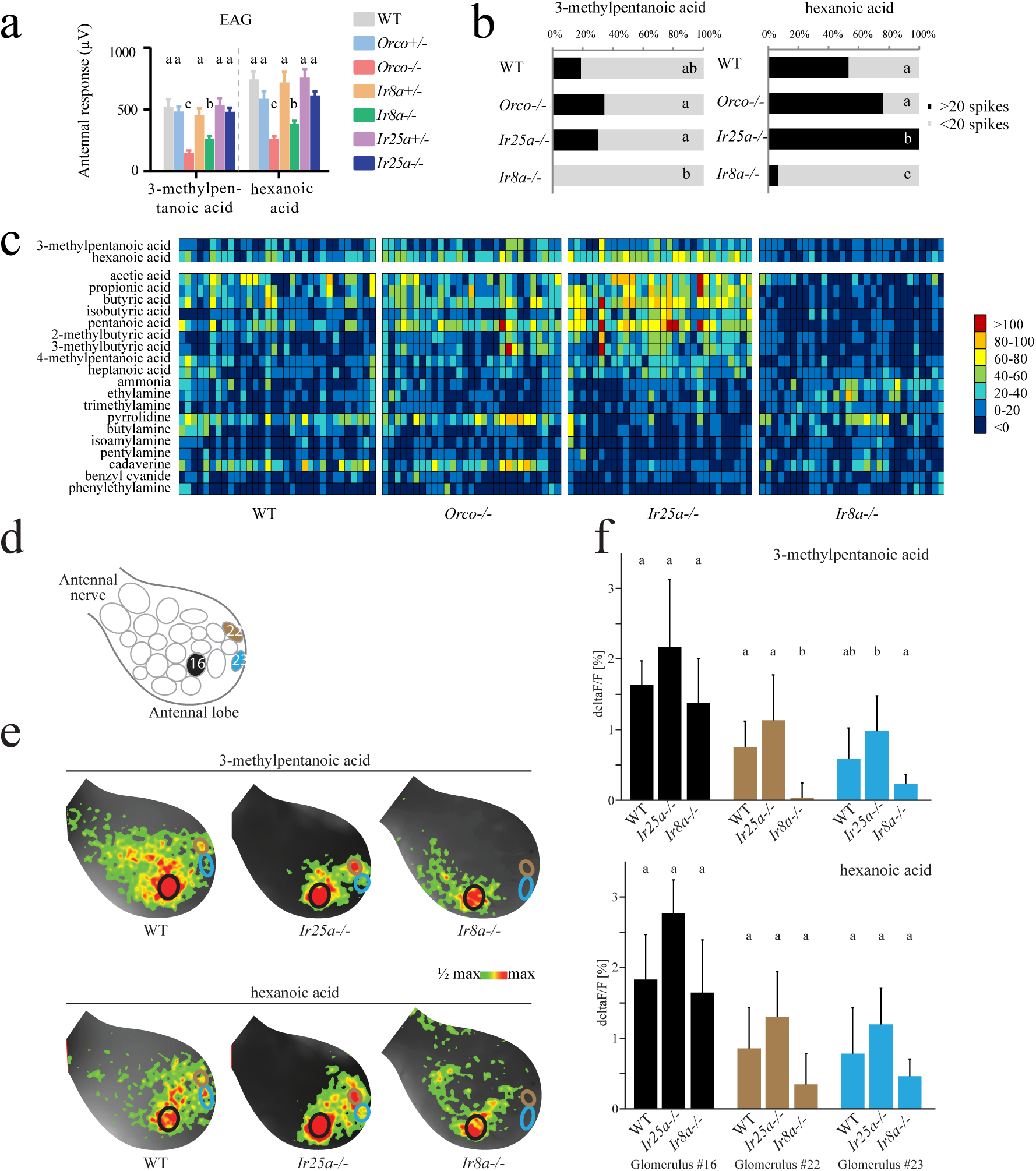
Detection and processing of frass-emitted. (**a**) Electroantennogram responses (EAG, in µV ± SEM, the response to solvent was subtracted) of *M. sexta* antennae isolated from wild type (WT), *Orco-/*- (Orco mutant), *Orco*+/− (Orco heterozygous), *Ir8a-/*- (Ir8a mutant), *Ir8a* +/− (Ir8a heterozygous), *Ir25a-/*- (Ir25a mutant), *Ir25a+/−* (Ir25a heterozygous). EAG responses to 3-methylpentanoic acid and hexanoic acid. (**b**) Percentage of coeloconic sensilla responding to the two behaviorally active acids in different genotypes (**c**) Heat map representation of SSR responses of coeloconic sensilla from different moth genotypes. (**d**) Schematic of 23 putative olfactory glomeruli at the dorsal surface of the right antennal lobe; the schematic was created for each individual moth based on the activation patterns of 19 diagnostic odorants^45^; numbers identify glomeruli that were most strongly activated by the tested acids (#16), or that showed acid-specific activation (#22, #23). (**e**) Examples of calcium imaging recordings in wildtype, *IR25a*−/−, and *IR8a*−/− female moths after stimulation with the two behaviorally active acids. The increase of fluorescence is color coded (see scale) and superimposed onto the view of the antennal lobe; circles indicate positions of glomeruli #16 (black outline), #22 (brown), #23 (blue). (**f**) Bars show the mean response of a glomerulus (after subtraction of the solvent response) to an odorant; error bars indicate standard deviation; bars with the same letter are not significantly different from each other (ANOVA, n=4-6 females/genotype).

**Figure 3.**
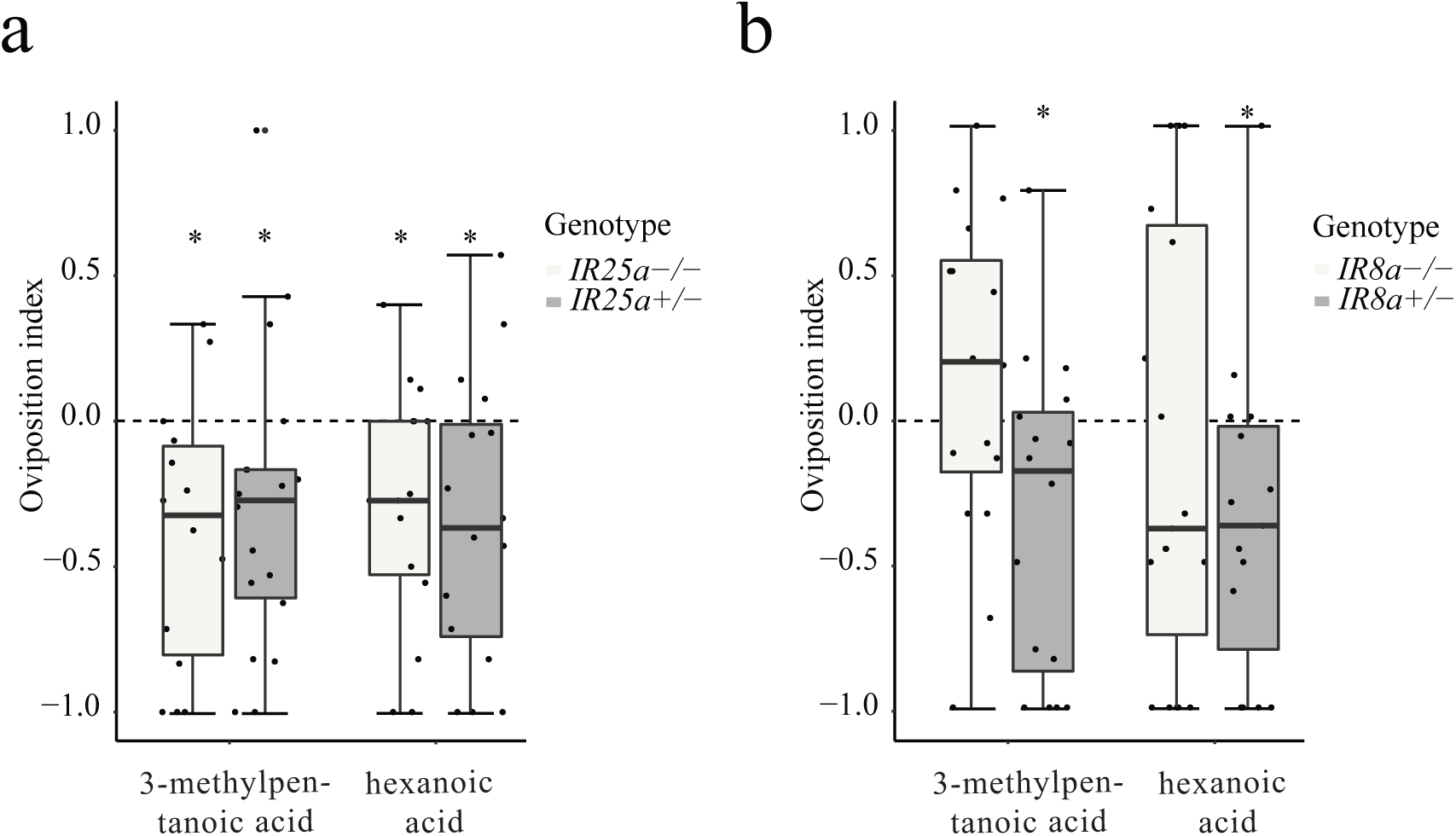
IR8a is necessary for acid avoidance of ovipositing *M. sexta* females. Two-choice assay showing the oviposition indexes of the homozygous and heterozygous (as a control) of *Ir25a* (**a)** and *Ir8a* (**b**) mutants for the frass-emitted compounds 3-methylpentanoic acid and hexanoic aicd (for details on choice assay see Fig. 1 and methods section).

To address whether the remaining response to acids in *Orco*−/− moths were indeed resulting from activation of the IR pathway, we generated two IR mutant lines, *Ir8a*−/− and *Ir25a*−/−, using again CRISPR/Cas9 genome editing. The resulting *Ir8a*−/− mutant contained a 339bp deletion (93bp at exon2, 170bp at intron2 and 76bp at exon3) while the *Ir25a*−/− mutant contained a 154bp deletion (154bp at exon2) in the genome. As both deletions resulted in frame-shifts and the occurrence of premature stop codons (Supplementary Fig. 4A), we expected both mutations to result in non-functional ionotropic co-receptors. We found no difference regarding pupal weight and length in neither *Ir8a*−/− nor *Ir25a*−/− mutants, when compared to the heterozygous controls (Supplementary Fig. 4B). Furthermore, in EAG experiments both mutants exhibited normal responses to the OR-detected pheromone, bombykal (Supplementary Fig. 4C) suggesting the absence of relevant off-target effects.

However, when performing EAG experiments with *Ir8a*−/− and *Ir25a*−/− moths, only *Ir8a*−/− moths exhibited significantly reduced response to both behaviorally active acids when compared to WT moths, while the acid responses in *Ir25a*−/− moths remained unaffected (Fig. 2a).

### IR8a pathway is essential for detecting and avoiding acids from caterpillar frass

We next asked which sensillum type is involved in the detection of the acids in caterpillar frass. According to the well-studied *Drosophila* ^9, 18, 39, 40^, IR-expressing OSNs are mainly housed in coeloconic sensilla. Furthermore, in *M. sexta*, previous single-sensillum recordings (SSRs) from trichoid and basiconic sensilla showed little to no responses to acids^41, 42^. We therefore hypothesized that coeloconic sensilla of *M. sexta* house IR-expressing OSNs that are involved in acid detection. Contrary to the antenna of female *D. melanogaster*, which contains only 54 coeloconic sensilla^40^, the antenna of female *M. sexta* carries about 3600^41^. This makes the identification and recording from identified individual coeloconic sensilla almost impossible. We, therefore, recorded from 28 coeloconic sensilla from the middle part of the antenna, which should cover a wide range of functional types, and stimulated them with a set of 52 odorants from different chemical classes (Supplementary Fig. 5). Consistent with previous studies in *D. melanogaster*^40^ and *Bombyx mori*^43^, OSNs housed in coeloconic sensilla in wild type *M. sexta* were mainly activated by acids and amines. The two behaviorally active acids activated mainly OSNs in non-overlapping groups of coeloconic sensilla. The number of coeloconic sensilla responding to hexanoic acid was about two times higher than those responding to 3-methylpentanoic acid and the intensity of responses to hexanoic acid was stronger.

We found reduced numbers of coeloconic sensilla responding to 3-methylpentanoic acid and hexanoic acid in *Ir8a*−/− moths, when comparing to the other three genotypes (Fig. 2b and c). Interestingly, increased numbers of coeloconic sensilla exhibited enhanced responses to acids in *Ir25a*−/− moths, whereas the responses to amines were almost abolished. The enhanced responses in *Ir25a-/*- moths toward acids could be due to more energy being available to OSNs responding to acids, as these OSNs no longer have to compete with amine-tuned OSNs in the same sensillum. Such a phenomenon has been reported for gustatory receptors (GRs), where sensory neurons in the same sensillum have been shown to interact, exhibiting competition, inhibition or activation^44^.—In conclusion, our results show that IR8a, but neither Orco nor IR25a, is required for acid detection in OSNs of coeloconic sensilla.

We conclude that IR8a is involved in the detection of the key compounds governing frass avoidance. We next asked, where in the antennal lobe (i.e. the first olfactory processing center of the moth’s brain) this IR-related acid detection becomes processed. A recently published functional analysis of the moths’ antennal lobe^45^ revealed three glomeruli that strongly responded to acids. In another study^30^, activation of two of these glomeruli was not affected by knocking out the OR-coreceptor Orco, supporting that these two glomeruli become innervated by IR-expressing OSNs. When performing calcium imaging experiments with moths that either lacked a functional IR25a or IR8a, the responses to acids in *Ir25a* mutants were unaffected compared to control animals (Fig. 2 e, f). However, when testing *Ir8a* mutants, we observed a slightly reduced response to hexanoic acid and a significantly reduced response to 3-methylpentanoic acid (Fig. 2 e, f) in only those two glomeruli that in the former study were independent of Orco. Together with the EAG results, we conclude that both Orco and IR8a, but not IR25a, are involved in acid sensing and that IR8a-expressing OSNs involved in the detection target a subset of glomeruli on the medial surface of the antennal lobe.

Finally, we asked whether any of the three co-receptors governs the behavioral avoidance towards acids in ovipositing *M. sexta*. Unfortunately, the oviposition rates of *Orco*−/− moths were too low to draw any conclusions regarding the involvement of Orco in the oviposition avoidance. One explanation for the conflicting result with our previews study^30^ in terms of oviposition in *Orco*−/− moths is that Fandino et al (2019) used whole and large *D. wrightii* plants which probably provided a very strong visual stimulation. Interestingly, however, mutation of *Ir25a* did not affect the oviposition behavior (Fig. 3a), while *Ir8a*−/− moths were no longer repelled by the tested acids (Fig. 3b). We conclude that ovipositing females rely on IR8a for detection of acids from caterpillar frass.

Several studies have shown that larval frass and its odors deter female oviposition^36^. Frass-emitted acids play a crucial role in oviposition avoidance in moth species like *Ostrinia* species^46^ and *Helicoverpa armigera*^47^. Moreover, it was shown that female parasitoid wasps, *Cotesia glomerata*, use acids emitted by host larvae as cues to locate their host^48^, and *M. sexta* caterpillar frass-emitted acids play a major role in attracting predators like ants^33^. Finally, one of the acids we identified in the caterpillar frass (hexanoic acid) has been shown to induce plant defenses against herbivores^49^. Obviously, carboxylic acids are potent signals for a gravid female to realize that at a given plant the female’s offspring might face conspecific competitors, parasitoids and predators, as well as an already induced plant defense. Therefore, our finding that ovipositing *M. sexta* females, like other moths, avoid emitted acids from larval frass is not unexpected. However, the neural and molecular mechanisms as well as the exact chemistry underlying this behavior remained elusive. In this study, we do not only show that *M. sexta* display oviposition aversion toward caterpillar frass, but also find that only the major volatile compounds (C_6_ carboxylic acids) emitted are aversive for gravid females. By testing mutant moths in which we knocked out different olfactory co-receptors, we show that the co-receptors IR8a and Orco, but not IR25a, participate in acid detection. We also find that IR8a is necessary for the acid avoidance behavior of gravid *M. sexta* females, which helps the moth to protect its offspring from conspecific competition.

It has been reported that *M. sexta* lay significantly less eggs on plants treated with herbivore-induced volatile organic compounds due to high predation rate on those treated plants^31^. Our finding that *M. sexta* avoids competition by sensing not only plant-emitted, but also frass-emitted compounds adds another layer of regulation to host choice in *M. sexta* to distinguish between healthy and damaged plants.

## Methods

### Insect rearing and plant material

All animals were reared at the Max Planck Institute for Chemical Ecology, Jena, Germany, as already described^13^. Briefly, eggs were collected from female *M. sexta* moths, which could freely oviposit on *D. wrightii* plants. Larvae used in the experiments were reared on artificial diet, under 16:8 h light: dark photo period with a relative humidity of 40% at 26°C. Naïve females were mated the second night after emergence and tested during the subsequent night. *M. sexta* frass was collected daily from fourth to fifth instar caterpillars which were raised on either *N. attenuata or D. wrightii*.

All plants were grown in a greenhouse as described^50^. Plants used for experiments were not yet flowering. Approximately 7 days before being used, plants were transferred into a climate chamber with the same settings as the moth flight cage (16:8 h light: dark photo period with a relative humidity of 40% at 26°C.).

### Chemical analysis

We identified the volatiles of caterpillars frass using SPME (Solid Phase Microextraction) coupled with GC-MS (Gas chromatography–mass spectrometry). One gram of frass from caterpillars raised on either *D. wrightii* or *N. attenuata* were put into a 500-mL plastic container. A circular filter paper (diameter: 12 mm Whatman, Sigma-Aldrich USA) loaded with 10 µL of diluted bromodecane (1:10^4^ in hexane) was used as an internal standard. Through a hole in the lid of the container, a SPME fiber (50 µm Divinylbenzene / Carboxen / Polydimethylsiloxane coating; Supelco) was exposed to the container headspace for 30 min at room temperature without agitation, and then introduced into the injector inlet for 2 min at 250°C in split-less mode. The compounds adsorbed on the fiber were then analyzed by GC-MS (Agilent 6890 GC & 5975C MS, Agilent, USA). After fiber insertion, the column temperature was maintained at 40°C for 2 min and then increased to 260°C at 15°C min^−1^, followed by a final stage of 5 min at 260°C. Compounds were identified by comparing mass spectra against synthetic standards and NIST 2.0 library matches. All of the synthetic odorants that were tested and confirmed were purchased from Sigma (www.sigmaaldrich.com) and were of the highest purity available.

### Behavioral experiments in the wind tunnel

To investigate the behavioral significance of *M. sexta* frass from caterpillars which had fed on *N. attenuata*, we performed two choice tests in a transparent wind tunnel (220 × 90 × 90 cm^3^) at 25 °C, 70% relative humidity, 0.3 lux illumination, and a wind speed of 40 cm/s. Two non-flowering *N. attenuata* of similar size were placed at the upwind end of the wind tunnel. An empty petri dish (control) or a petri dish loaded with 10 gram of freshly collected frass (treatment) was placed at the base of the plant. A single 5^th^ instar larva produce about 10 gram of frass per day. As described before^45^, mated female moths were released at the downwind side of the wind tunnel and during 3 min were allowed to oviposit on both plants. Afterwards, the number of eggs on both plants was counted and the eggs were gently removed after each test. Moths were tested only once and plants were exchanged after two tests. The positions of treatment and control plant within the wind tunnel were swapped after every second moth. The oviposition indexes were calculated as (*T-C*)/(*T+C*) where *T* is the number of eggs on the treatment site and *C* is the number of eggs on the control site.

To test the effect of *M. sexta* frass from caterpillars that had raised on *D. wrightii*, we conducted a similar two choice test in the wind tunnel. Due to the large size of *Datura* plants, we trimmed plants seven days before the experiments in a way that two leaves of similar size remained on opposite directions. An empty petri dish (control) or a petri dish loaded with 10 gram of freshly collected frass (treatment) was placed 10 cm beneath the leaves. Again, mated female moths were allowed to oviposit on both control and treatment leaves and the resulting eggs and oviposition indexes were calculated afterwards.

To determine the functional significance of different volatiles emitted by the frass, we conducted two-choice tests in the wind tunnel. This time two freshly detached leaves of similar size were presented to the gravid female. Each leaf was attached to the tip of one of two upright acrylic glass poles (40 cm high and placed at the upwind end of the wind tunnel with a distance of 40 cm between them). Beneath each leaf we attached a square filter paper (2 × 2 cm^2^) loaded with 10 µL of diluted odorant (1:10^2^) or the solvent mineral oil alone. Moths, leaves and filter papers were tested only once. Experiments were conducted both with leaves from *N. attenuata* and *D. wrightii*.

### CRISPR/ Cas 9-based genome editing

To determine which co-receptor is involved in the acid detection and acid-driven oviposition avoidance, we used olfactory receptor co-receptor (Orco) mutant moths^30^, and generated Ionotropic receptor 8a (Ir8a) and Ir25a two mutant lines. The *M. sexta* genome v.1.0 (Mansexv1.0) fasta file and the GFF3 file were submitted to the CHOPCHOP (http://chopchop.cbu.uib.no) database for CRISPR/ Cas9 target selection sites. The OGS2.0 gene names, i.e. Msex2.10447-RB, isoform 1 and Msex2.02645-RA, isoform 1, were used to select target site. The sgRNA and Cas9 were synthesized by IDT (https://eu.idtdna.com/pages/products/crispr-genome-editing/alt-r-crispr-cas9-system). The microinjection and genotyping were carried out according to previously established procedures^30^. After the mutant lines were established, mutations were reconfirmed by Sanger sequencing.

### Electrophysiology

To investigate the antennal responses to frass-emitted carboxylic acids, we performed EAG (Electroantennography) recording. We therefore clipped the antenna of a 3-day-old female moth directly above the scapulum and before the third last flagellum. Antenna preparation, stimuli delivery, data acquisition and analysis were carried out according to previously established procedures^50^. Odorants for EAG analyses were selected based on compounds identified in the headspace of caterpillar frass as well as structurally similar chemicals. 10 µl of diluted odor (1:10^2^) or solvent alone were pipetted onto a circular filter paper (diameter: 12 mm) and placed into a glass pipette. In addition, we performed single-sensillum recordings from coeloconic sensilla as described before^42^. Coeloconic sensilla were identified by their characteristic morphology. 29-32 Coeloconic sensilla were recorded in each genotype. Responses were quantified by counting all spikes recorded from an individual sensillum due to difficulties in reliably distinguishing spikes from individual neurons^22, 40^. The response was calculated as the difference in spike number observed 0.5 s before and after the stimulus onset. Heatmap was generated in Excel. Calcium imaging experiments were conducted as described previously^45^. CAS number and purities for odorants is listed in supplementary table 1.

### Statistics and figure preparation

Sample size of behavioral experiments was determined based on a previous study^45^. Data were analyzed and plotted using RStudio (Version 1.1.414), R (Version 3.4.2; The R Project for Statistical Computing) and GraphPad InStat 3 (https://www.graphpad.com/scientific-software/instat/), while figures were organized and prepared using Adobe Illustrator CS5. The Wilks– Shapiro test was used to determine normality of each data set. Normally distributed data were assessed using t-tests. Not normally distributed data were analyzed using Wilcoxon signed-rank test, with the null hypothesis that the median of sampled values differs from zero. For the boxplots the whiskers were calculated as follows: the upper whisker equals the third quartile plus 1.5× the interquartile range (IQR) and the lower whisker equals the first quartile minus 1.5× the IQR. Any data points above the superior or below the inferior whisker values are considered as outliers. All data were included in the statistical analysis.

---

**Supplementary Figure 1.**
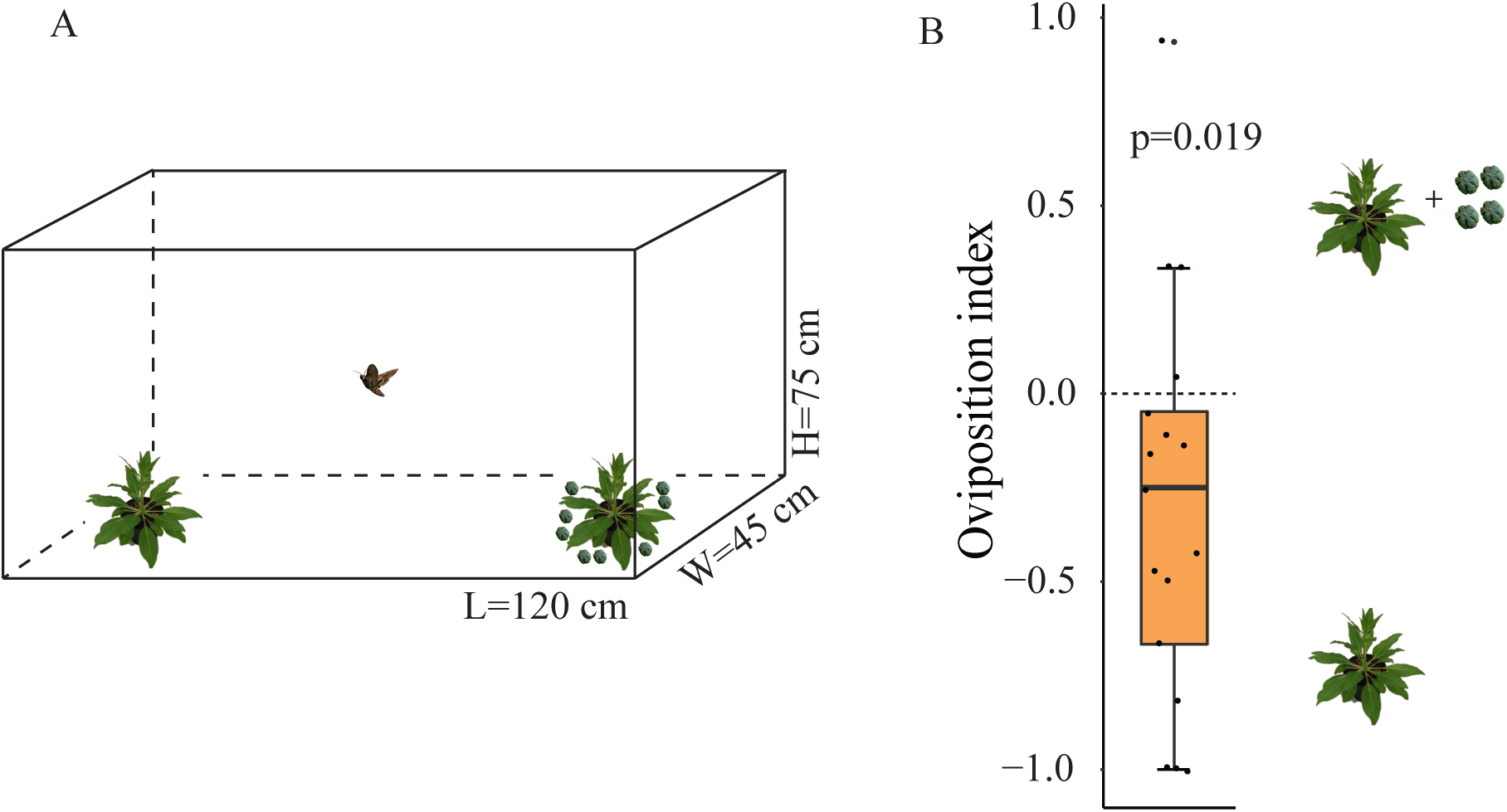
Frass avoidance of ovipositing *M. sexta* females is conserved in different behavioral assays. (A) Schematic drawing of oviposition cage. (B) Oviposition index of WT *M. sexta* given a choice between *N. attenuata* and *N. attenuata* containing caterpillar frass in the oviposition cage. Deviation of the index against zero was tested with Wilcoxon signed-rank test (n=20). * p<0.05. Boxplots depict median and upper and lower quartile; whiskers depict quartiles +/− 1.5× the interquartile range (IQR). Any data points above the superior or below the inferior whisker values are considered as outliers. All data were included in the statistical analysis.

**Supplementary Figure 2.**
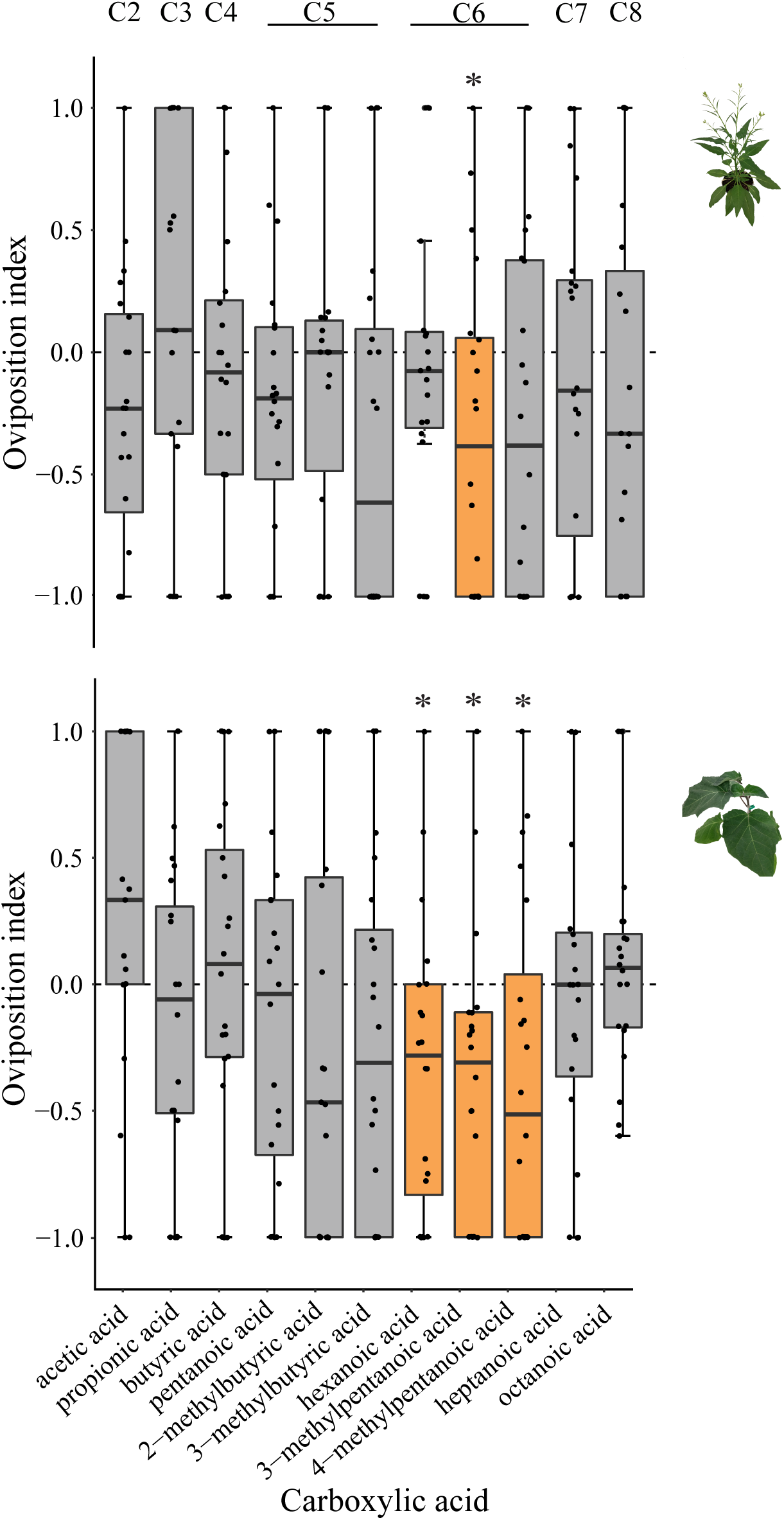
Host-plant-specific effects of different carboxylic acids on *M. sexta*’s oviposition choice. (**A**) Two-choice assay showing the preference of wild-type (WT) females for carboxylic acids over control in the context of detached *N. attenuata* leaf. (**B**) Two-choice assay showing the preference of wild-type (WT) females for carboxylic acids over control in the context of detached *D. wrightii* leaf. Deviation of the index against zero was tested with Wilcoxon signed-rank test (n=20). * p<0.05. Part of these data are already shown in Fig. 1 f. Please also consider that hexanoic acid obviously deters oviposition only in the right context, i.e. when females oviposit on *Datura wrightii*.

**Supplementary Figure 3.**
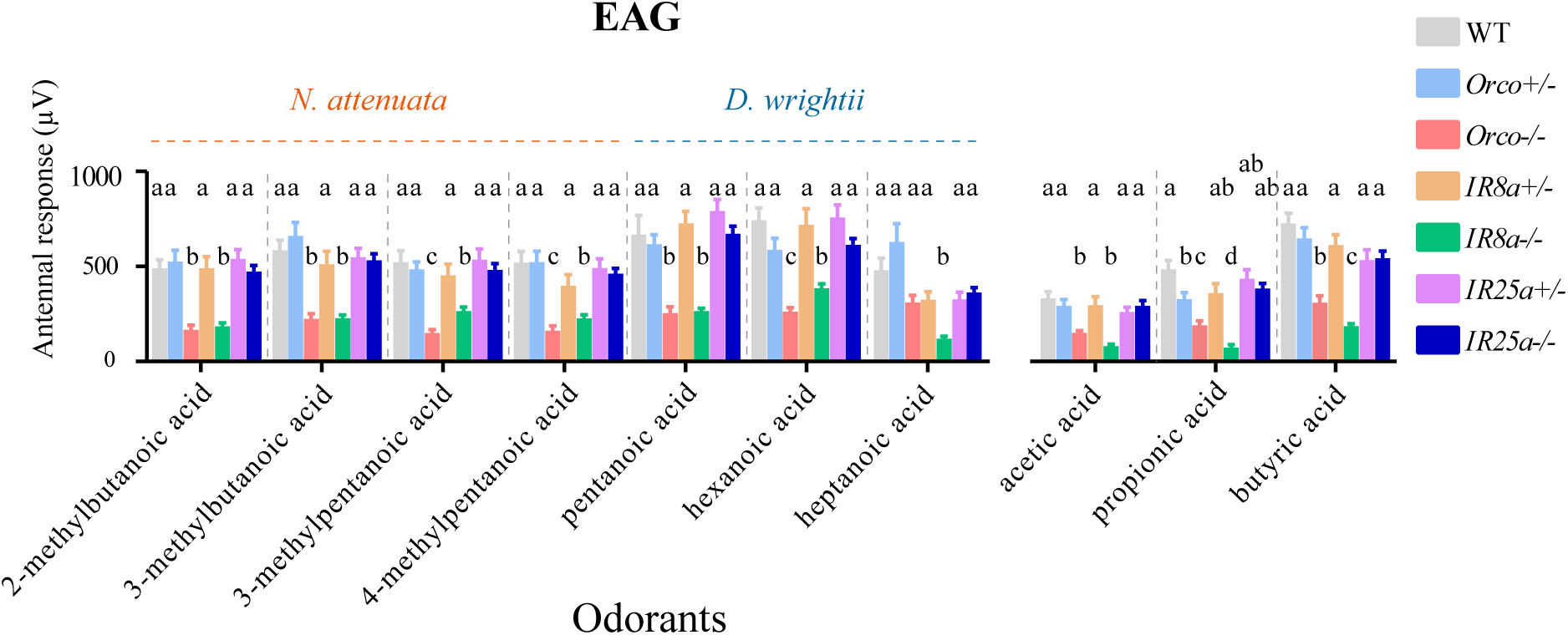
Both O*rco*−/− and *Ir8a*−/− *M. sexta* exhibit reduced electrophysiological responses to carboxylic acids. Electroantennogram responses (EAG, in mV ± SEM, the response to solvent was subtracted) of *M. sexta* antennae isolated from wild-type (WT), O*rco*−/− (*Orco* mutant), *Orco*+/− (*Orco* heterozygous), *Ir8a*−/− (*Ir8a* mutant), *Ir8a* +/− (*Ir8a* heterozygous), *Ir25a*−/− (*Ir25a* mutant), *Ir25a* +/− (*Ir25a* heterozygous). Responses to 3-methylpentanoic acid and hexanoic acid have been shown in Figure 2. Different letters above each odorant response indicate significant differences (one-way anova; p < 0.05); a is different from b and c, and b is different from c and ab is not different from either a or b.

**Supplementary Figure 4.**
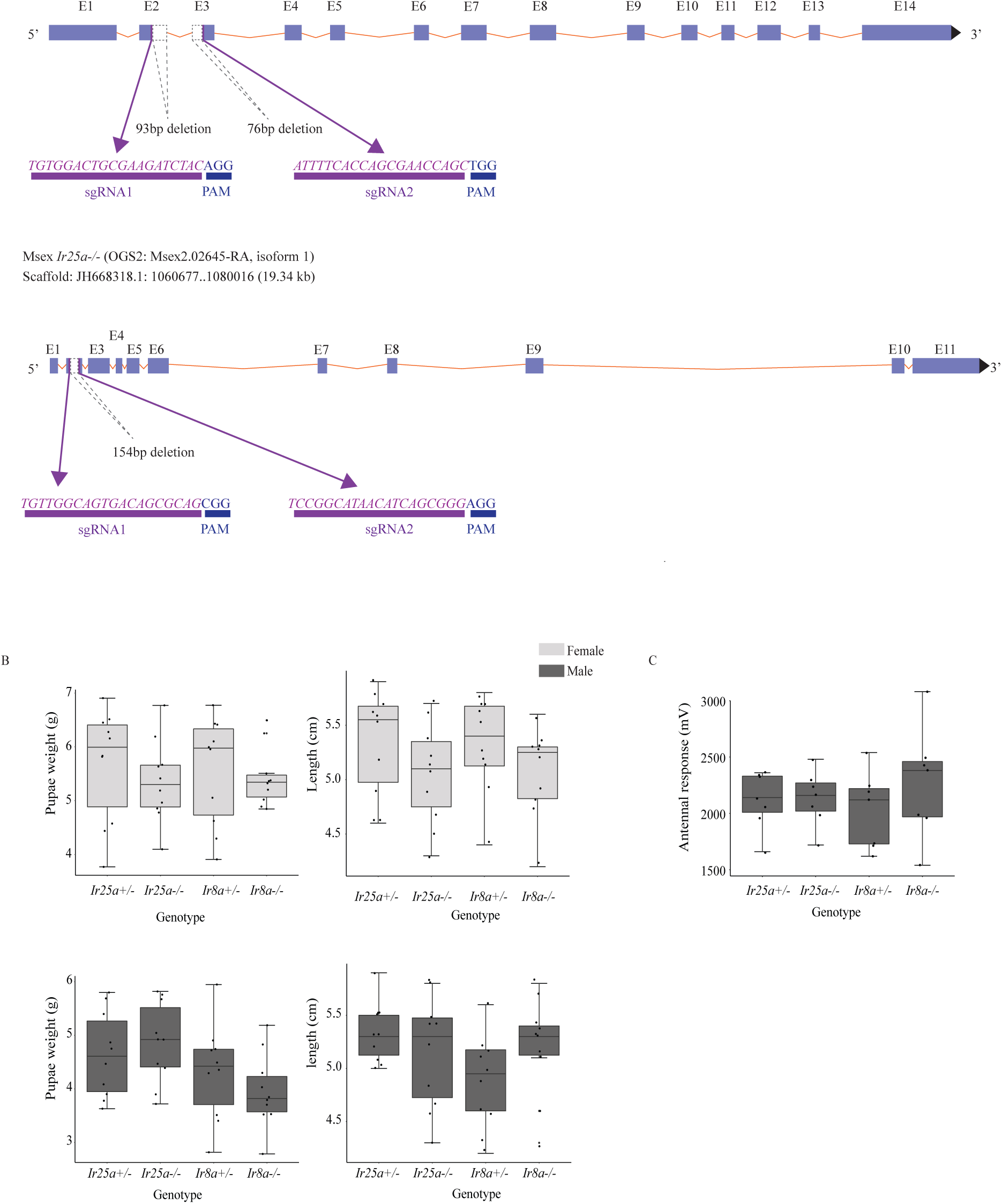
Gene editing technique and potential off-target effects. (**A**) **Schematic of the genes targeted via CRISPR-Cas9.** *Ir8a*−/− mutant contained a 339bp deletion (93bp at exon2, 170bp at intron2 and 76bp at exon3) while the *Ir25a*−/− mutant contained a 154bp deletion (154bp at exon2) in the genome. PAM protospacer adjacent motif. (B) Weight (left) and length (right) of female pupae from both *Ir8a* (upper panels) and *Ir25a* (lower panels) mutant and heterozygous lines. There were no statistical differences among corresponding genotypes (Wilcoxon signed-rank test, n=10). (C) EAG response of male adults toward pheromone (bombykal) in all genotypes. There were no statistical differences among genotypes (one-way ANOVA, n=7).

**Supplementary Figure 5.**
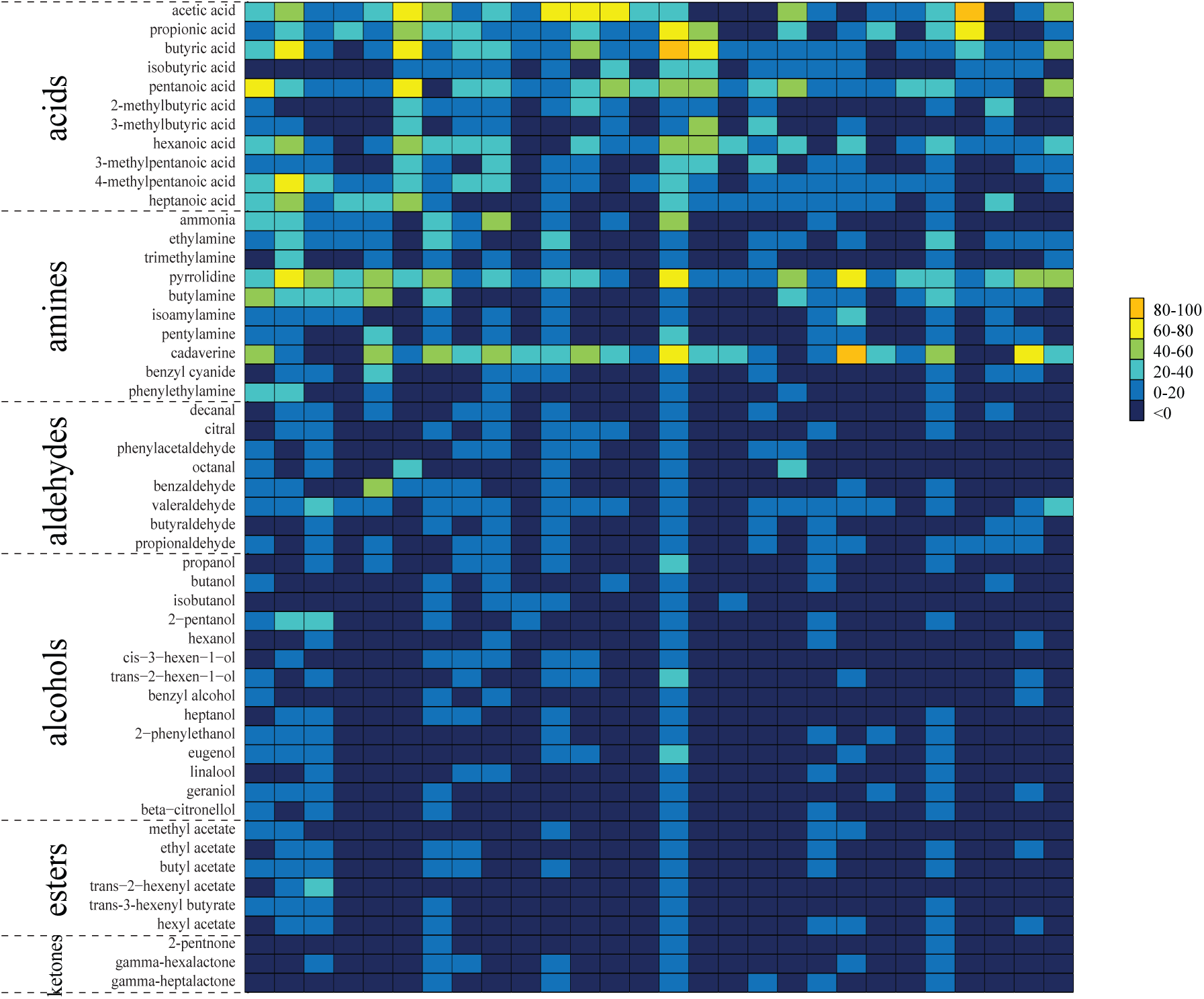
Heatmap based on SSRs of 28 coeloconic sensilla from WT *M. sexta* to 52 screened odors. Color-coded numbers depict difference in spikes/0.5s before and after stimulus onset.

**Supplementary Figure 6.**
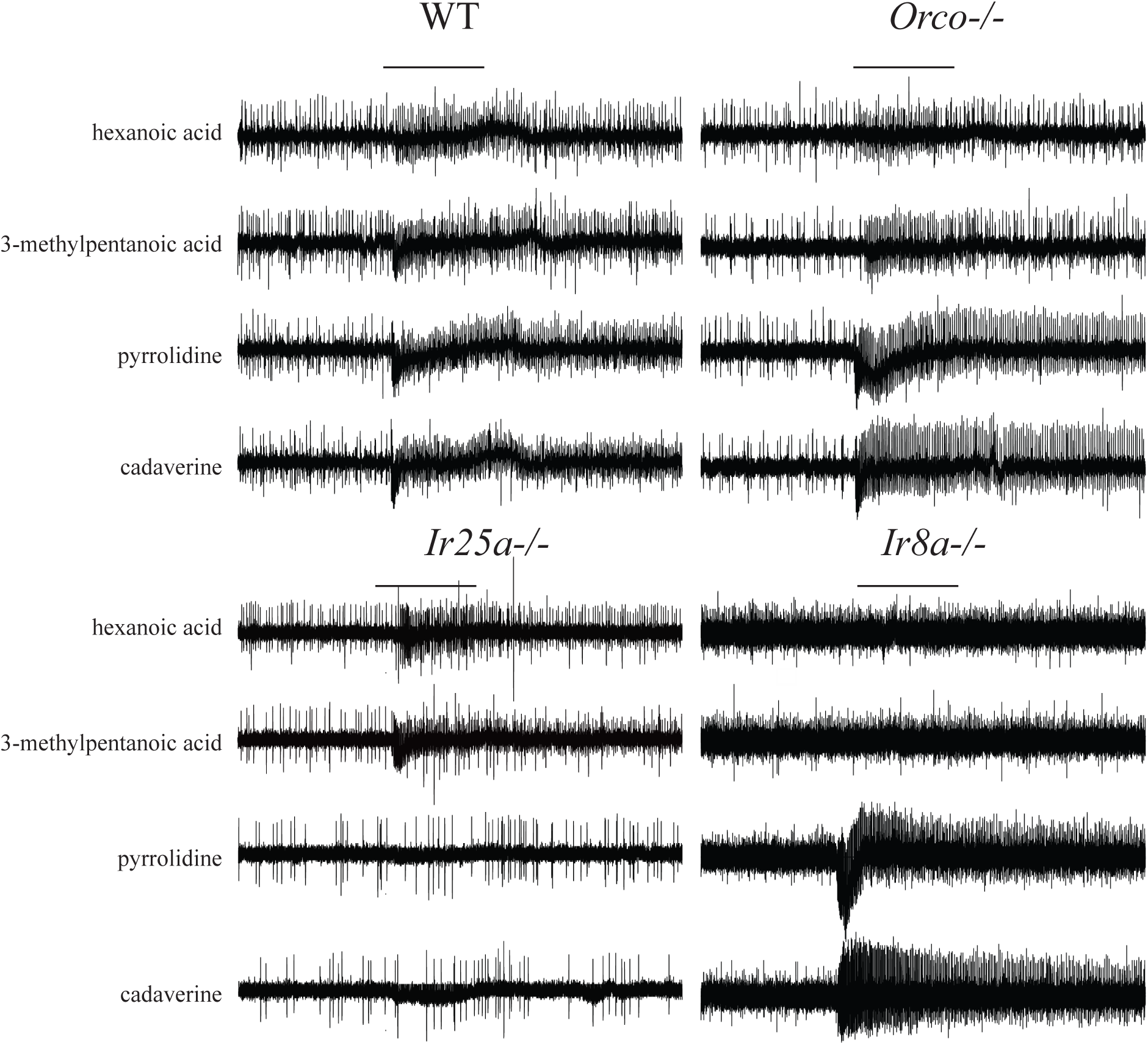
Representative SSR traces of coeloconic sensilla. Bars above the traces mark 0.5 s stimulus time.

## Acknowledgements

We thank the glasshouse team of Max Planck Institute for Chemical Ecology for plant cultivation; Sascha Bucks for rearing *M. sexta*; Vajiheh Jafari, Dima Ward, Syed Ali Komail Raza for help with the wind tunnel experiments. We express our gratitude to Kerstin Weniger for her technical support. This work was supported by the Max Planck Society (B.S.H) and the Alexander von Humboldt Foundation (J.Z.).

## Author contributions

J.Z., M.K. and B.S.H. designed the study; J.Z. performed all SSR and GC-MS experiments. J.Z. and S.W.Y. conducted wind tunnel assays, EAG experiments and measured pupae weight. J.Z., R.F., G.F.O., and E.G.W designed sgRNA and established CRISPR/Cas9 knockouts. S.B.K. performed the calcium imaging experiments. The original manuscript was written by J.Z., subsequently edited by M.K., S.B.K. and B.S.H., and all coauthors contributed to the final version of this paper.

---

**Table S1.** List of 52 tested stimuli.

| Odorant name | CAS Number | Odorant name | CAS Number |
| --- | --- | --- | --- |
| acetic acid | 64-19-7 | propanol | 71-23-8 |
| propanoic acid | 79-09-4 | butanol | 71-36-3 |
| butyric acid | 107-92-6 | isobutanol | 78-83-1 |
| isobutyric acid | 79-31-2 | 2-pentanol | 6032-29-7 |
| pentanoic acid | 109-52-4 | hexanol | 111-27-3 |
| 2-methylbutyric acid | 116-53-0 | cis-3-hexen-1-ol | 928-96-1 |
| 3-methylbutyric acid | 503-74-2 | trans-2-hexen-1-ol | 928-95-0 |
| hexanoic acid | 142-62-1 | benzyl alcohol | 202-859-9 |
| 3-methylpentanoic acid | 105-43-1 | heptanol | 111-70-6 |
| 4-methylpentanoic acid | 646-07-1 | 2-phenylethanol | 60-12-8 |
| heptanoic acid | 111-14-8 | eugenol | 97-53-0 |
| ammonia | 7664-41-7 | linalool | 78-70-6 |
| ethylamine | 75-04-7 | geraniol | 106-24-1 |
| trimethylamine | 75-50-3 | beta-citronellol | 106-22-9 |
| pyrrolidine | 123-75-1 | methyl acetate | 79-20-9 |
| butylamine | 109-73-9 | ethyl acetate | 141-78-6 |
| isoamylamine | 107-85-7 | butyl acetate | 123-86-4 |
| pentylamine | 110-58-7 | trans-2-hexenyl acetate | 2497-18-9 |
| cadaverine | 462-94-2 | trans-3-hexenyl butyrate | 53398-84-8 |
| benzyl cyanide | 140-29-4 | hexyl acetate | 142-92-7 |
| phenylethylamine | 64-04-0 | 2-pentnone | 107-87-9 |
| decanal | 112-31-2 | gamma-hexalactone | 695-06-7 |
| citral | 5392-40-5 | gamma-heptalactone | 105-21-5 |
| phenylacetaldehyde | 122-78-1 | octanal | 124-13-0 |
| butyraldehyde | 123-72-8 | benzaldehyde | 100-52-7 |
| propionaldehyde | 123-38-6 | valeraldehyde | 110-62-3 |

